# Embryo-like features in developing *Bacillus subtilis* biofilms

**DOI:** 10.1101/2020.03.09.983718

**Authors:** Momir Futo, Luka Opašić, Sara Koska, Nina Čorak, Tin Široki, Vaishnavi Ravikumar, Annika Thorsell, Domagoj Kifer, Mirjana Domazet-Lošo, Kristian Vlahoviček, Ivan Mijaković, Tomislav Domazet-Lošo

**Affiliations:** Laboratory of Evolutionary Genetics, Ruđer Bošković Institute, Bijenička cesta 54, HR-10000 Zagreb, Croatia; Department for Evolutionary Theory, Max Planck Institute for Evolutionary Biology, August-Thienemann-Str. 2, D-24306 Plön, Germany; Faculty of Electrical Engineering and Computing, University of Zagreb, Unska 3, HR-10000 Zagreb, Croatia; The Novo Nordisk Foundation Center for Biosustainability, Technical University of Denmark, 2800 Kgs. Lyngby, Denmark; Proteomics Core Facility, Sahlgrenska Academy, University of Gothenburg, Medicinaregatan 5, SE-41390 Gothenburg, Sweden; Faculty of Pharmacy and Biochemistry, University of Zagreb, A. Kovačića 1, HR-10000 Zagreb, Croatia; Bioinformatics Group, Division of Biology, Faculty of Science, University of Zagreb, Horvatovac 102a, HR-10000 Zagreb, Croatia; University of Skövde, School of Biosciences, Högskolevägen, Box 408, 541 28 Skövde, Sweden; Systems and Synthetic Biology Division, Department of Biology and Biological Engineering, Chalmers University of Technology, Kemivägen 10, SE-41296 Gothenburg, Sweden; Catholic University of Croatia, Ilica 242, HR-10000 Zagreb, Croatia

## Abstract

Correspondence between evolution and development has been discussed for more than two centuries^1^. Recent work reveals that phylogeny-ontogeny correlations are indeed present in developmental transcriptomes of eukaryotic clades with complex multicellularity^2–10^. Nevertheless, it has been largely ignored that the pervasive presence of phylogeny-ontogeny correlations is a hallmark of development in eukaryotes^6, 10–12^. This perspective opens a possibility to look for similar parallelisms in biological settings where developmental logic and multicellular complexity are more obscure^13–16^. For instance, it has been increasingly recognized that multicellular behaviour underlies biofilm formation in bacteria^13, 14, 17–19^. However, it remains unclear whether bacterial biofilm growth shares some basic principles with development in complex eukaryotes^14–16, 18, 20^. Here we show that the ontogeny of growing Bacillus subtilis biofilms recapitulates phylogeny at the expression level. Using time-resolved transcriptome and proteome profiles, we found that biofilm ontogeny correlates with the evolutionary measures, in a way that evolutionary younger and more diverged genes were increasingly expressed towards later timepoints of biofilm growth. Molecular and morphological signatures also revealed that biofilm growth is highly regulated and organized into discrete ontogenetic stages, analogous to those of eukaryotic embryos^11, 21^. Together, this suggests that biofilm formation in Bacillus is a bona fide developmental process comparable to organismal development in animals, plants and fungi. Given that most cells on Earth reside in the form of biofilms^22^ and that biofilms represent the oldest known fossils^23^, we anticipate that the widely-adopted vision of the first life as a single-cell and free-living organism needs rethinking.

---

Multicellular behaviour is wide-spread in bacteria and it was proposed that they should be considered multicellular organisms^13^. However, this idea has not been generally adopted likely due to the widespread laboratory use of domesticated bacterial models selected against multicellular behaviours, the long tradition of viewing early diverging groups as simple, and the lack of evidence for system-level commonalities between bacteria and multicellular eukaryotes^15, 16, 18, 24^. Recently developed phylo-transcriptomic tools for tracking evolutionary signatures in animal development^2–4^ were also successfully applied in the analysis of developmental processes in plants and fungi^5, 6, 10^. Although development evolved independently in these three major branches of eukaryotic diversity^12^, their ontogenies showed similar phylogeny-ontogeny correlations indicating that possibly all eukaryotic developmental programs have an evolutionary imprint. Transferability of the phylo-transcriptomic tools across clades and likely universal patterns of phylogeny-ontogeny correlations in eukaryotic multicellularity prompted us to test this approach on bacterial biofilms − a multicellular, and the most common, form of bacterial existence in nature^22^.

*Bacillus subtilis* NCIB3610, a frequently used model organism in bacterial biofilm research^17^, shows many properties readily found in multicellular eukaryotes including cell differentiation, division of labour, cell signalling, morphogenesis, programmed cell death and self-recognition^17–19^. Nevertheless, these properties are often seen as collective behaviours of independent cells rather than features of a multicellular individual. Here we altered the perspective by approaching the biofilm as an individual organism and applying phylo-transcriptomics as it would be used for studying ontogeny of an embryo-forming eukaryote.

## Biofilm growth is a stage-organised process

To measure transcriptome expression levels during *B. subtilis* biofilm formation, we sampled eleven timepoints covering a full span of biofilm ontogeny from its inoculation, until two months of age (Fig. 1a). We recovered transcriptome expression values for 4,316 (96%) *B. subtilis* genes by RNAseq, which revealed three distinct periods of biofilm ontogeny: early (6H-1D), mid (3D-7D) and late period (1M-2M), linked by two transition stages at 2D and 14D (Fig. 1b, c, Extended Data Fig. 1, Supplementary Table 1). Biofilm transcriptomes showed a time-resolved principal component analysis (PCA) profile (Extended Data Fig. 1) and poor correlation to the liquid culture (LC) used for inoculation of biofilms (Fig. 1b), indicating that biofilm makes a distinct part of the *B. subtilis* life cycle. When we considered all ontogeny timepoints, 4,266 (99%) genes were differentially expressed. This number stayed similar (4,199, 97% genes) when we looked only at biofilm growth *sensu stricto* (6H-14D, Supplementary Video 1) by excluding the starting liquid culture (LC) and late-period timepoints (1M-2M) that show biofilm growth arrest. These values reflect highly dynamic regulation of transcription in biofilm ontogeny, comparable to those seen in animal embryos^2, 11^. Pairwise comparisons between successive ontogeny timepoints uncovered that most genes (around 70%) change their transcription at biofilm inoculation (LC-6H), indicating that transition from a liquid culture to solid agar represents a dramatic shift in *Bacillus* lifestyle (Extended Data Fig. 2). The two most dynamic steps during biofilm growth are transitions at 1D-2D and 7D-14D where, respectively, around 30% and 25% *B. subtilis* genes change transcription (Extended Data Fig. 2). Together with correlation (Fig. 1b), PCA (Extended Data Fig. 1) and clustering analyses (Fig. 1c, Supplementary Information 1), this shows that biofilm growth is not a continuous process^14^. Instead, like development in animals^11, 21^, it is punctuated with bursts of transcriptional change that define discrete ontogeny phases (the early, mid and late period).

**Fig. 1.**
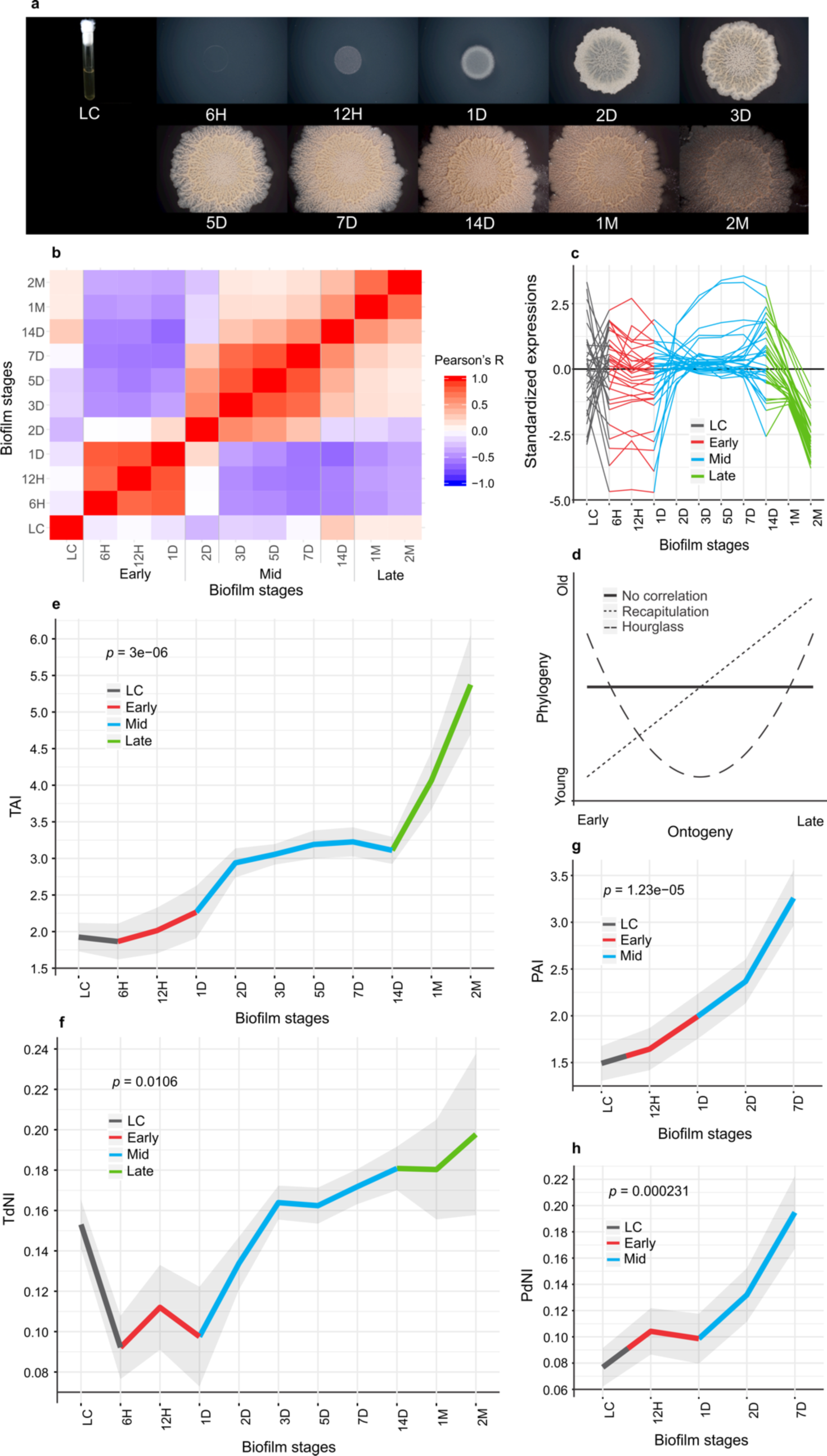
*B. subtilis* biofilm growth is a highly regulated and punctuated process exhibiting a phylogeny-ontogeny recapitulation pattern. a, Gross morphology of *B. subtilis* biofilms on solid agar plates at 6 hours (6H), 12 hours (12H), 1 day (1D), 2 days (2D), 3 days (3D), 5 days (5D), 7 days (7D), 14 days (14D), 1 month (1M) and 2 months (2M) after inoculation with liquid culture (LC) (see Supplementary Video 1). **b**, Pearson’s correlation coefficients between timepoints of biofilm ontogeny in all-against-all comparison. Early (6H-1D), mid (3D-7D) and late (1M-2M) periods, together with transition stages at 2D and 14D, are marked. **c**, Average transcriptome expression profiles of 31 the most populated gene clusters (for all 64 clusters see Supplementary Information 1). **d**, Hypothetical profiles of phylogeny-ontogeny correlations. Solid line displays no correlation, dotted line the recapitulation model and dashed line the hourglass model. **e**, Transcriptome age index (TAI) and **g,** proteome age index (PAI) profiles of *B. subtilis* biofilm ontogeny show recapitulation pattern. **f**, Transcriptome nonsynonymous divergence index (TdNI) and **h**, proteome nonsynonymous divergence index (PdNI) profiles show that genes conserved at nonsynonymous sites are used early in the biofilm ontogeny, while more divergent ones later during biofilm ontogeny. Nonsynonymous divergence rates were estimated in *B. subtilis* - *B. licheniformis* comparison. Depicted *p* values are obtained by the flat line test and grey shaded areas represent ± c estimated by permutation analysis (see Methods, **e-h**). Early (red), mid (blue) and late (green) periods of biofilm growth are colour coded (**c**, **e-h**).

**Fig. 2.**
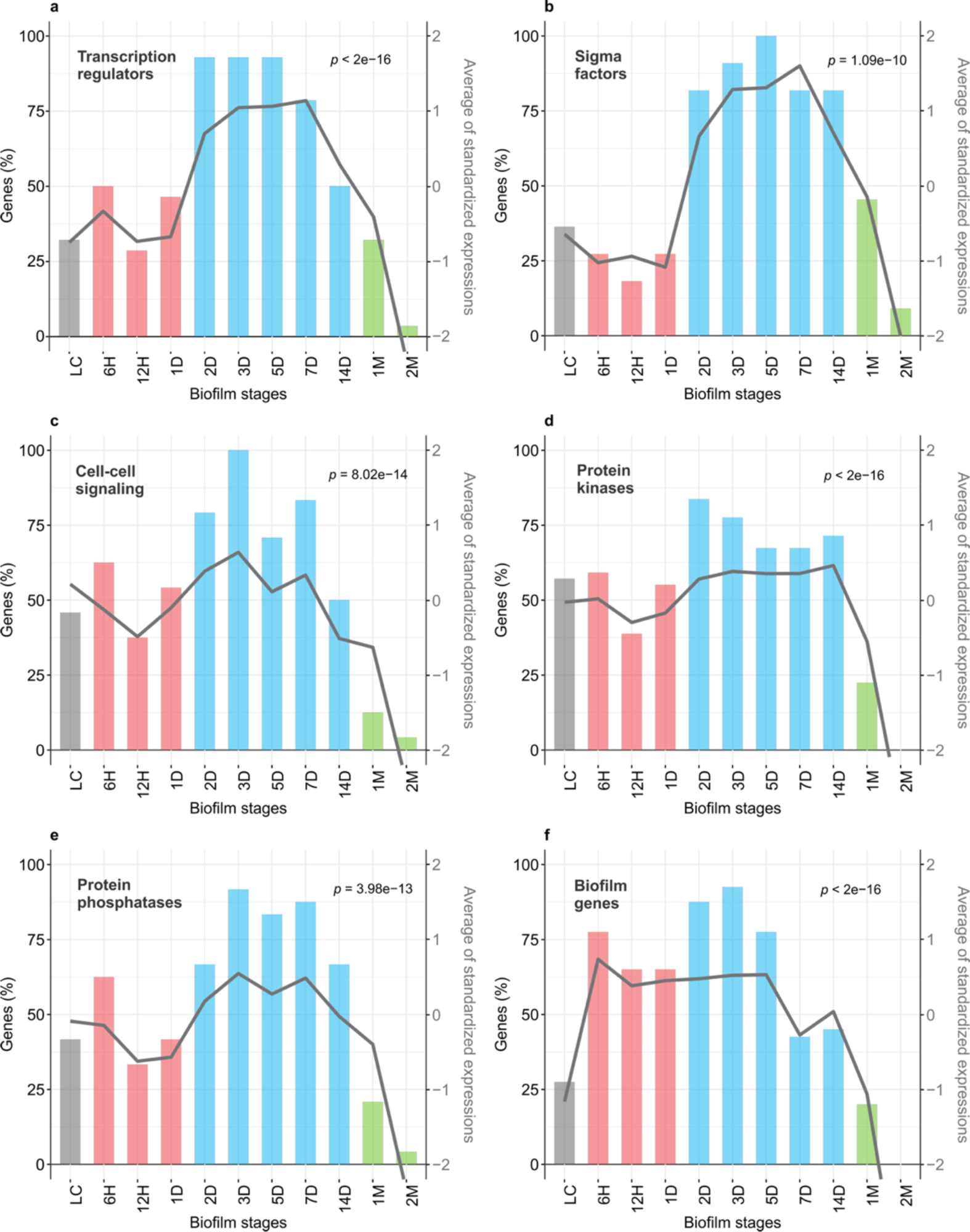
Multicellularity-important genes show cumulatively the strongest transcription in the mid-biofilm period. Left y-axis shows percentage of genes that are transcribed above the median of their overall transcription profile (histogram). Right y-axis shows the average standardized transcription values for all considered genes (line). Significance of the average expression profile is tested by repeated measures ANOVA and respective *p* values are shown. **a**, Transcription regulators that regulate ≥ 10 operons (see Methods, n = 28, *F*(10, 270) = 17.33); **b**, Sigma factors (n = 11, *F*(10, 100) = 9.257); **c**, Cell to cell signalling genes (n = 24, *F*(10, 230) = 9.947); **d**, Protein kinases (n = 49, *F*(10, 480) = 41.71); **e**, Protein phosphatases (n = 24, *F*(10, 230) = 9.452); **f**, Key biofilm genes (n = 40, *F*(10, 390) = 30.74). Colouring of bars in histograms follows biofilm growth periods: LC (grey), early (red), mid (blue), late (green). See Extended Data Fig. 7 and Extended Data Fig. 9 for full profiles of all considered genes.

## Evolutionary expression measures show a recapitulation pattern

To assess whether biofilm growth has some evolutionary directionality, or if this process is macroevolutionary naive, we linked transcriptome profiles to evolutionary gene age estimates to obtain the transcriptome age index (TAI); a cumulative measure that gives overall evolutionary age of an expressed mRNA pool^2, 5, 6^. Evolutionary gene age estimates are obtained by phylostratigraphic procedure^25^ using consensus phylogeny (Extended Data Fig. 3, Supplementary Information 2, Supplementary Table 2). If one assumes that expression patterns across biofilm ontogeny are independent of evolutionary age of genes, then the TAI profile should show a trend close to a flat line; i.e. TAI and ontogeny should not correlate (Fig. 1d). Alternatively, there are many possible phylogeny-ontogeny correlation scenarios that would reflect a macroevolutionary imprint, but hourglass and recapitulation pattern are the two mostly considered (Fig. 1d). Although the recapitulation pattern historically was the first one to be proposed^1^, recent studies of eukaryotic development mainly support the hourglass model^2–7, 9, 10^. Surprisingly, in *B. subtilis* we found a recapitulation pattern where early timepoints of biofilm growth express evolutionary older transcriptomes compared to mid and late timepoints that exhibit increasingly younger transcriptomes (Fig. 1e). This correlation between biofilm timepoints (ontogeny) and TAI (phylogeny) indicates that, like in complex eukaryotes, a macroevolutionary logic plays a role in *B. subtilis* biofilm formation. We also examined how the TAI profile relates to the evolutionary age of genes (phylostrata - ps) and found that recapitulation pattern is significant already from the origin of Firmicutes (ps4; Extended Data Fig. 4), reflecting its rather deep roots in the bacterial phylogeny (Extended Data Fig. 3).

**Fig. 3.**
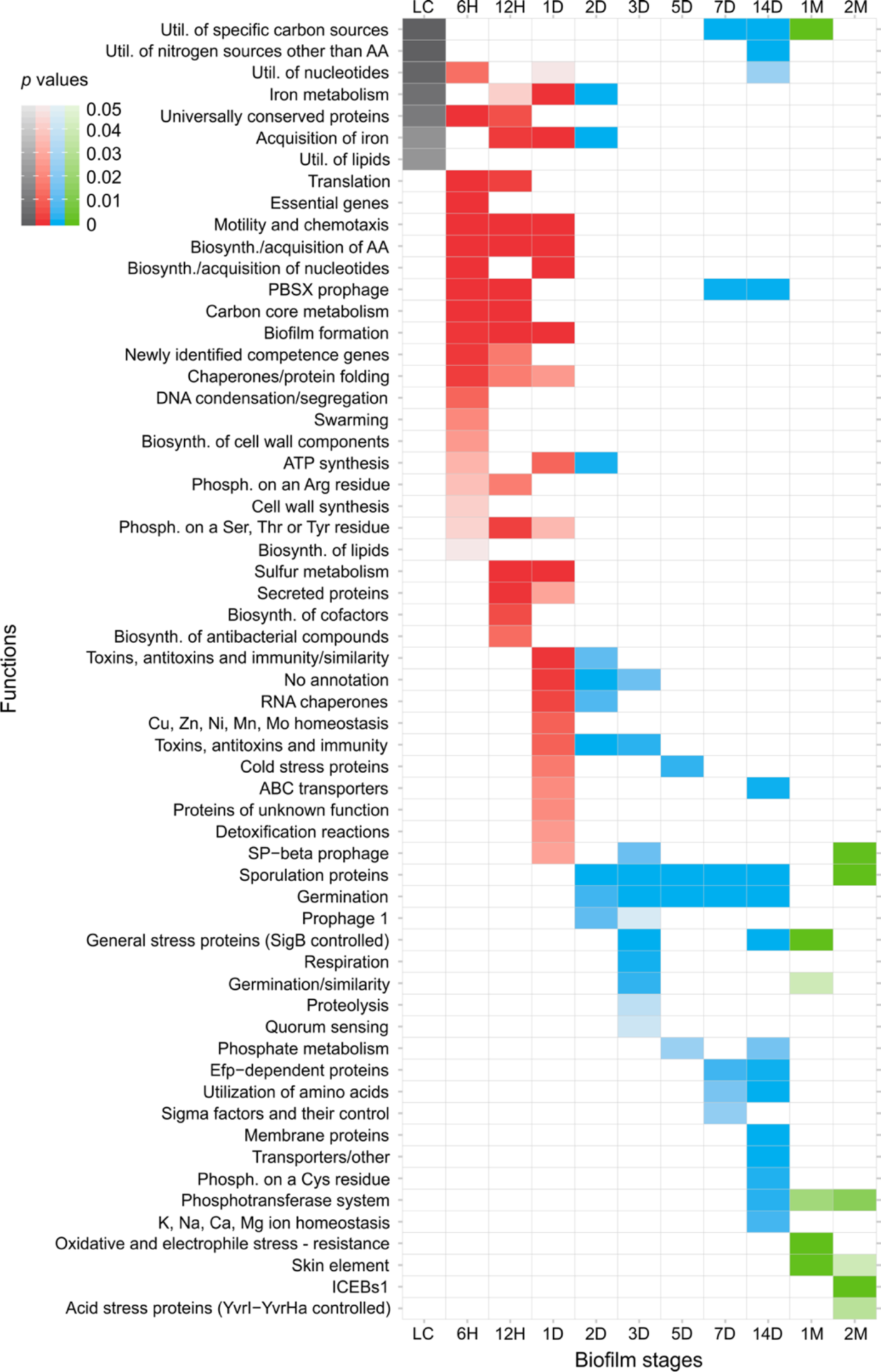
Biofilm ontogeny is a punctuated process organized in functionally discreate stages. Enrichment analysis of SubtiWiki functional categories (ontology depth 3) in a respective biofilm growth timepoint for genes with transcription 0.5 times (log2 scale) above the median of their overall transcription profile. Similar results are obtained for other transcription level cut-offs and SubtiWiki functional annotation ontology depths (see Supplementary Table 4). Colouring follows biofilm growth periods: LC (grey), early (red), mid (blue), late (green). Functional enrichment is tested by one-tailed hypergeometric test and *p* values are adjusted for multiple testing (see Methods).

Phylostratigraphic procedure used in determining gene ages is based on detecting remote homologs^2, 25^ (Extended Data Fig. 3, Supplementary Table 2). However, one could also analyse the dataset by looking at more recent evolutionary history via estimating evolutionary divergence rates of coding sequences^5^. It is usually assumed that nonsynonymous substitution rates (dN) reflect selective pressures, in contrast to synonymous substitution rates (dS) that provide an estimate of neutral evolution in coding sequences. However, due to the strong codon usage bias, selection also acts on synonymous sites in *B. subtilis*, and therefore its dS rates cannot be considered neutral^26^. To account for this, we looked at substitution rates separately by devising transcriptome nonsynonymous (TdNI) and synonymous (TdSI) divergence indices (see Methods). In *B. subtilis - B. licheniformis* comparison, from 1D onwards TdNI showed a recapitulation pattern where genes conserved at nonsynonymous sites tend to be used early, while more divergent ones are used later during the biofilm ontogeny (Fig. 1f, Supplementary Table 3). Comparably, TdSI displays more complex correlation which clearly resembles the pattern of the transcriptome codon bias index (TCBI) indicating dependence of synonymous substitution rates and codon usage bias (Extended Data Fig. 5). Nevertheless, TdSI recapitulation profile is evident in mid-period biofilms (1D-14D), where genes with more divergent synonymous sites gradually increase in transcription from 1D to 14D (Extended Data Fig. 5). Together, these divergence-ontogeny parallelisms in *B. subtilis* biofilms further corroborate the recapitulative evolutionary imprint and show that it is actively maintained by relatively recent evolutionary forces in mid-period biofilms.

Regulated mRNA transcription is the essential step in gene expression. However, a full molecular phenotype visible to selection is reached only after protein translation. In non-steady state processes like ontogeny, a plethora of opposing factors influence mRNA and protein levels, resulting in their relatively low correlations^27^. To test if the recapitulation pattern also exists at the proteome level, we quantified proteomes of representative stages (LC, 12H, 1D, 2D, 7D), which cover the most dynamic part of macroscopic morphology change (Supplementary Video 1). We obtained protein expression values for 2,907 (67%) predicted proteins (Supplementary Table 1) and used them to calculate the proteome age index (PAI); a cumulative measure analogous to TAI (see Methods), that gives an overall evolutionary age of a protein pool. Regardless of the relatively poor correspondence between transcriptome and proteome levels within timepoints (Extended Data Fig. 6a-g), and across ontogeny (Extended Data Fig. 6h, i), the PAI profile also showed a significant recapitulation pattern where evolutionary older proteins have higher expression early and younger ones later during biofilm ontogeny (Fig. 1g). Similar to TdNI and TdSI for transcriptome, proteome nonsynonymous (PdNI; Fig. 1h) and synonymous (PdSI; Extended Data Fig. 5) divergence indices in *B. subtilis-B. licheniformis* comparison revealed that recapitulation pattern also holds at shallower evolutionary levels (see Methods). Jointly, this demonstrates that phylogeny-ontogeny dependence, beside transcriptomes, is also visible in biofilm proteomes.

## Multicellularity important genes dominate in mid-period biofilms

To find further parallels between biofilm growth and multicellular development, we looked for expression patterns of transcription factor and cell-cell signalling genes, which are defining features of development in complex eukaryotes^28^. We found that *Bacillus* transcription regulators cumulatively have the highest transcription in mid-period biofilms (2D to 7D) and that during this period almost all of them are transcribed above the median of their overall expression profiles (Fig. 2a). This holds even if we narrow down the analysis to sigma factors only (Fig. 2b), and mimics developing transcriptomes in animals where embryos show increased expression of transcription factors^28^. Similarly, quorum sensing genes peak in transcription at 3D (Fig. 2c), suggesting the most elaborate cell-cell communication at the timepoint when the biofilm gets the typical wrinkled morphology (Fig. 1a; Supplementary Video 1). Protein phosphorylation is another important mechanism involved in cell signalling and differentiation both in eukaryotes and bacteria, and it plays a critical role in *B. subtilis* biofilm formation^17–19^. Again, we found that protein phosphorylation genes (kinases and phosphatases) cumulatively have the highest transcription in mid-period biofilms (Fig. 2d, e), likely reflecting various types of cell differentiation in this growth phase^18, 19^.

Functional annotation of *B. subtilis* genes, including the gene regulatory networks underlying biofilm formation^17–19^, is comparably advanced in terms of quality and completenes^29^. This allowed us to follow the expression of key biofilm genes and analyse specific functional patterns in biofilm ontogeny (Fig. 2f, Fig. 3). Collectively, the key biofilm genes are increasingly transcribed from the onset of biofilm formation (6H), maintain high values over early and mid-period, and progressively decline in late biofilms (Fig. 2f). Their individual profiles, however, reflect their specific roles. For instance, extracellular matrix genes show highest transcription in early biofilms (6H-1D), sporulation and cannibalism genes have highest transcription in the mid-period (2D-14D), surfactin has bimodal distribution with peaks at 6H and 14D and protease production increases from 2D to 14D (Extended Data Fig. 7, Supplementary Information 3 and 4).

## Biofilm growth has a stepwise functional architecture

The functional category enrichment analysis of biofilm timepoints reveals a tight control where every timepoint expresses a specific battery of functions (Fig. 3, Extended Data Fig. 8, Supplementary Table 4). Some illustrative examples of enriched functions include PBSX prophage (eDNA production), antibacterial compounds and swarming in early biofilms (6H-12H), zinc metabolism at 1D, iron uptake siderophores and functionally unannotated genes at transition stage 2D, quorum sensing at 3D, sporulation and toxins/antitoxins in the mid-period (2D-14D), general stress at transition stage 14D, and mobile genetic elements in late biofilms (1M-2M). The enrichment of genes that lack functional annotation at 2D probably reflects the incomplete knowledge on the molecular mechanisms that govern early-to-mid biofilm transition. Statistical analysis of these genes on the phylostratigraphic map (Extended Data Fig. 10) reveals that they preferentially originate from the ancestors of *B. subtilis* strains (ps10-ps12). This parallels development in animals where phylogenetically restricted genes (orphans) are involved in the embryonic transitions and the generation of morphological diversity^2, 30, 31^. When observed in total, functional enrichment patterns show that biofilm growth at the functional level has discreate hierarchical organization with even finer temporal grading compared to the pure transcription profiles (Fig. 1b, 3, Extended Data Fig. 8). This modular nature of biofilm growth is analogous to the non-continuous and stage-organized architecture of development in animals^11, 14, 21^.

## Conclusions

Bonner proposed that polarity and heritable pattern formation, including cell differentiation with division of labour, are fundamental properties of any development^32^. *B. subtilis* biofilms indeed show polarity along bottom-top^17, 18^ and central-distal axes^33^, along with remarkably complex cell differentiation with division of labor^17, 18, 34^. Long range electrical signaling^35^, control of cheater cells^19^ and recent discovery of cancer-like processes in aging biofilms^36^ are additional features that go far beyond these minimal conditions that define multicellular development. In this study we showed that phylogeny-ontogeny correlations and stage-organized gene expression architecture should be added to the list of properties that qualify *B. subtilis* biofilm growth as a true multicellular developmental process, analogous to developmental processes in complex eukaryotes. It is somewhat surprising that the recapitulation pattern originally proposed to be present in animals by 19th century zoologists^1^ is now actually found in bacteria. However, this proves that the cross-talk between zoology and microbiology could bring new and exciting insights^37^. For example, in the light of multicellular individuality of *B. subtilis* biofilms, a host-bacterial symbiosis that involves biofilms^17, 38^ could be viewed as an interaction of the two multicellular individuals.

Multicellularity is not a rare evolutionary transition as it has independently evolved many times in various lineages^12, 15^. However, at every independent occurrence, it seems to be governed by the similar basic principles that include a macroevolutionary imprint. Future work should establish generality of these findings across bacterial and archaeal diversity as well as ecological conditions including microbial community biofilms. Yet, the results of our study, pervasiveness of bacterial^22^ and archaeal multicellular behavior^39^, and the fact that the first fossils were bacterial biofilms^23^, encourage us to call for the re-evaluation of the widely adopted idea that the first life on the Earth was unicellular. It is undisputable that the cell is the basic unit of life; however, that does not readily imply that the first life was strictly unicellular. At least some models envisage that protocells were organized in biofilm-like structures^40^, and that unicellular part of the life cycle could evolve in parallel as an efficient dispersion mechanism in early oceans^20, 41^.

## Contributions

T.D-L. initiated and conceptualized the study; T.D-L. and I.M. supervised the study; L.O. and N.Č. optimized protocols for culturing bacteria, RNA and proteome isolation; L.O. and N.Č. isolated transcriptomes and proteomes; M.F., K.V. and S.K. processed raw transcriptome data, mapped reads and establish expression levels; I.M., V.R. and A.T. performed LC-MS experiments and quantified proteomes; M.F., K.V., S.K., T.Š. and T.D-L. analysed the data and prepared the figures; M.D-L. developed phylostratigraphic pipeline; N.Č., M.D-L., D.K. and T.D-L. constructed consensus phylogeny and built the database; M.F., S.K., N.Č. and M.D-L. made phylostratigraphic maps; T.D.-L. and M.F. wrote the manuscript with contributions of all authors. All authors read and approved the manuscript.

## Acknowledgements

We thank Ž. Pezer-Sakač, P. Štancl, B. Pavletić and I. Šutevski for assistance with protein isolation. T.C.G. Bosch, A. Klimovich and G. Klobučar for discussions. This work was supported by the Novo Nordisk Foundation (NNF10CC1016517) to I.M., and City of Zagreb Grant, Croatian Science Foundation under the project IP-2016-06-5924, Adris Foundation Grant and European Regional Development Fund Grants KK01.1.1.01.0008 (CERRM) and KK.01.1.1.01.0009 (DATACROSS) to T.D-L.

## Competing interests

The authors declare no competing interests.

## Methods

### Biofilm growth

*Bacillus subtilis* subsp. *subtilis* str. NCIB3610 (*B. subtilis*) was obtained from the Bacillus Genetic Stock Center (BGSC, Ohio State University, Columbus, OH, USA) and stored in 25% glycerol stocks at -80 °C. Bacteria from the stock were plated on a LB agar plate (1% Bacto tryptone, 0.5% Bacto yeast extract, 1% NaCl, 1 mM NaOH solidified with 1.5% agar) and incubated for 24h at 37 °C. Liquid LB medium (10 mL) were inoculated with a single colony and incubated with shaking for 24h at 37 °C and 250 rpm. Petri dishes (90 mm) containing MSgg agar (5 mM potassium phosphate pH 7, 100 mM MOPS pH 7, 2 mM MgCl2, 700 µM CaCl2, 50 µM MnCl2, 50 µM FeCl3, 1 µM ZnCl2, 2 µM thiamine, 0.5% glycerol, 0.5% glutamate, 50 µg/mL tryptophan, 50 µg/mL phenylalanine solidified with 1.5% agar) were inoculated with four drops (5 μL) of LB culture. The drops on each plate were approximately equidistantly distributed. The plates were incubated at 30 °C and the biofilms were harvested for RNA extraction at 6 and 12 hours, and at 1, 2, 3, 5, 7, 14, 30 and 60 days post-inoculation time (transcriptome samples 6H, 12H, 1D, 2D, 3D, 5D, 7D, 14D, 1M and 2M, respectively). For protein extraction biofilms were harvested at 12 hours and at 1, 2, and 7 days post-inoculation time (proteome samples 12H, 1D, 2D and 7D, respectively).

### RNA extraction

To reach a satisfactory amount of biomass for RNA extraction, 102 (6H), 34 (12H) and four (1D, 2D, 3D, 5D, 7D, 14D, 1M and 2M) biofilms were pooled per sample. The starting liquid LB culture (LC) used for biofilm inoculation was pelleted by centrifugation. All samples, excluding 2M, were taken in three biological replicates per timepoint. We succeeded to get only one replicate for 2M due to technically demanding RNA extraction from aged biofilms. To prevent changes in RNA composition due to biofilm harvesting procedure, 1 mL of stabilization mix (RNAprotect Bacteria Reagent - Qiagen diluted with PBS in a 2:1 volume ratio) was applied on plates. Soaked biofilms were gently removed from the agar surface using a sterile Drigalski spatula and a pipette tip and together with the unabsorbed stabilization mix transferred into a 2 mL tube (Eppendorf). An additional 1 mL of stabilization buffer was added to the tubes and the content was homogenized with a sterile pestle. The total RNA was extracted by applying a modified version of the RNeasy Protect Mini Kit (Qiagen) protocol. The homogenized samples were vortexed for 10 sec, resuspended by pipetting and incubated for 5 min at RT. 300 μL of the homogenate were transferred into a new 1.5 mL tube (Eppendorf), centrifuged for 10 min at 5000 x *g* at RT and 220 μL of the mix containing 200 μL of TE buffer, 3 mg of lysozyme and 20 μL of Proteinase K were added. The tubes were incubated in a shaker for 30 min at 25 °C and 550 rpm. 700 μL of RLT buffer were added into tubes, vortexed for 10 sec, and the suspension was transferred into a 2 mL tube containing 50 mg of 425 – 600 μm acid-washed glass beads (Sigma-Aldrich). The bacterial cells were disrupted using a Fast prep FP120 homogenizer (Thermo Savant Bio101) at 6.5 m/sec shaking speed for 5 min. After homogenization, the tubes were centrifuged for 15 sec at 13,000 x *g* and 760 μL of the supernatant were transferred into a new 2 mL tube. 300 μL of chloroform were added, followed by a vigorous shaking by hand for 15 sec. After incubation for 10 min at RT, the tubes were centrifuged for 15 min at 13,000 x *g* and 4 °C. The upper phase was gently removed into new 1.5 mL tubes and 590 μL of 80% ethanol were added. The suspension was gently mixed by pipetting and 700 μL of it was transferred into a mini spin column and centrifuged for 15 sec at 13,000 x *g*. This step was repeated until the total volume of suspension was filtered through the mini spin column. The columns were washed in three steps using 700 μL of RW1 buffer (Qiagen) and two times 500 μL or RPE buffer (Qiagen) with centrifugation for 15 sec at 13,000 x *g*. After the last washing step, the columns were centrifuged for 2 min at 13,000 x *g*. The columns were transferred into new 1.5 mL tubes. 50 μL of RNase-Free water (Qiagen) were applied directly on the column filter and incubated for 1 min at RT. The columns were centrifuged for 1 min at 13,000 x *g* and discarded afterwards. 10 μL of the RDD buffer (Qiagen), 2.5 μL of DNase I (Qiagen) and 37.5 μL of RNase-Free water were added to the filtrate. After 10 min of incubation at RT, 50 μL of 7.5 M LiCl solution were added. The tubes were incubated for 1h at -20 °C. After incubation, the tubes were centrifuged for 15 min at 13,000 x *g* and 4 °C, the supernatant was discarded and 150 μL of 80% ethanol were added. After another centrifugation step for 15 min at 13,000 x *g* and 4 °C, the supernatant was discarded, and samples were resuspended in 30 μL of RNase-Free water. The RNA quantification was performed using a NanoDrop 2000 spectrophotometer (ThermoFisher Scientific). The RNA was stored at -80 °C until sequencing.

### RNAseq

Ribosomal RNA was removed from the total RNA samples by Ribo-Zero rRNA Removal Kit (Bacteria) (Illumina). RNA-seq libraries were prepared using the Illumina TruSeq RNA Sample Preparation v2 Kit (Illumina). The RNA sequencing was performed bi-directionally on the Illumina NextSeq 500 platform at the EMBL Genomics Core Facility (Heidelberg, Germany), generating ∼450 million reads per run. Before mapping, the sequence quality and read coverage were checked using FastQC V0.11.7^42^ with satisfactory outcome for each of the samples. In total 1,493,163,656 paired-end sequences (75bp) were mapped onto the *B. subtilis* reference genome (NCBI Assembly accession: ASM205596v1; GCA_002055965.1) using BBMap V37.66^43^ with an average of 93.46% mapped reads per sample. The *SAMtools* package V1.6^44^ was used to generate, sort and index BAM files for downstream data analysis. Subsequent RNAseq data processing was performed in R V3.4.2^45^ using custom-made scripts. Briefly, mapped reads were quantified per each *B. subtilis* open reading frame using the R *rsamtools* package V1.30.0^46^ and raw counts for 4,515 open reading frames were retrieved using the *GenomicAlignments* R package V1.14.2^47^. Expression similarity across timepoints and replicates was assessed using principal component analysis implemented in the R package *DESeq2* V1.18.1^48^ and visualized using custom-made scripts based on the R package *ggplot2* V3.1.0^49^ (Extended Data Fig. 1).

### Protein digestion

To reach a satisfactory amount of biomass for protein digestion, four biofilms were pooled per sample (12H, 1D, 2D and 7D). The LC sample were obtained by pelleting starting liquid LB culture. The samples were taken in three replicates at each timepoint. After corresponding incubation periods, 1 mL of cell lysis buffer (4% w/v SDS, 100 mM TEAB pH 8.6, 5 mM β-Glycerophosphate, 5 mM NaF, 5 mM Na3VO4, 10 mM EDTA and 1/10 tablet of Mini EDTA-free Protease Inhibitor Cocktail, Sigma-Aldrich) was used to harvest biofilms from the plates. Soaked biofilms were gently removed from the agar surface using a sterile Drigalski spatula or a pipette tip and together with the unabsorbed lysis buffer were transferred into a 2 mL tube. The bacterial biomass was resuspended in the cell lysis buffer and boiled for 10 min at 90 °C. After boiling, the samples were sonicated on ice for 30 min at 40% amplitude using the homogenizer Ultrasonic processor (Cole Parmer), and finally centrifuged for 30 min at 13,000 x *g* and 4 °C. The supernatant was discarded and the pellet was cleaned by chloroform/methanol precipitation (4 vol. of 99.99% methanol, 1 vol. of chloroform and 3 vol. of milliQ water). The lysate was centrifuged for 10 min at 5,000 x *g* and 4 °C. The upper, aqueous phase was discarded without disturbing the interphase. Four volumes of methanol were added to the tube, vortexed and centrifuged for 10 min at 5,000 x *g* and 4 °C. The supernatant was discarded and the pellet was air-dried for 1 min. The air-dried pellet was dissolved in a denaturation buffer (8 M Urea, 2 M Thiourea in 10 mM Tris-HCl pH 8.0). Samples were separated on a NuPage Bis-Tris 4–12% gradient gel (Invitrogen) based on the manufacturer’s instructions. 100 µg of total protein at each timepoint was loaded on the gel and run for a short duration. The gel was stained with Coomassie blue and subsequently cut into three slices (fractions). Resulting gel slices were destained by washing thrice with 5 mM ammonium bicarbonate (ABC) and 50% acetonitrile (ACN). Gel pieces were next dehydrated in 100% ACN. Proteins were then reduced with 10 mM Dithiothreitol in 20 mM ABC for 45 min at 56

°C and alkylated with 55 mM iodoacetamide in 20 mM ABC for 30 min at RT in dark. This was followed by two more washes with 5 mM ABC and 50% ACN and once with 100 % ACN. Proteins were digested with Trypsin protease (Pierce^TM^) at 37 °C overnight. Resulting peptides were extracted from the gel in three successive steps using the following solutions - Step 1: 3% trifluoroacetic acid in 30% ACN; Step 2: 0.5% acetic acid in 80% ACN; Step 3: 100% ACN. Extracted peptides were next concentrated in a vacuum centrifuge and desalted using C18 stage-tips^50^. Briefly, C18 discs (Empore) were activated with 100% methanol and equilibrated with 2% ACN, 1% TFA. Samples were loaded onto the membrane and washed with 0.5% acetic acid. Peptides were eluted in 80% ACN, 0.5% acetic acid, concentrated in a vacuum centrifuge, and analysed on the mass spectrometer.

### Mass Spectrometry

The MS analyses were performed at The Proteomics Core Facility at the Sahlgrenska Academy (University of Gothenburg, Sweden). Samples were analysed on an QExactive HF mass spectrometer interfaced with Easy-nLC1200 liquid chromatography system (Thermo Fisher Scientific). Peptides were trapped on an Acclaim Pepmap 100 C18 trap column (100 µm x 2 cm, particle size 5 µm, Thermo Fischer Scientific) and separated using an in-house constructed C18 analytical column (300 x 0.075 mm I.D., 3 µm, Reprosil-Pur C18, Dr. Maisch) using the gradient from 6% to 38% acetonitrile in 0.2% formic acid over 45 min followed by an increase to 80% acetonitrile in 0.2% formic acid for 5 min at a flow of 300 nL/min. The instrument was operated in data-dependent mode where the precursor ion mass spectra were acquired at a resolution of 60,000, the ten most intense ions were isolated in a 1.2 Da isolation window and fragmented using collision energy HCD settings at 28. MS2 spectra were recorded at a resolution of 15,000 (for the 12H and 1D timepoints) or 30,000 (for the 2D, 7D and LC timepoints), charge states two to four were selected for fragmentation and dynamic exclusion was set to 20 s and 10 ppm. Triplicate injections (technical replicates) were carried out for each of the sample fractions for label free quantitation (LFQ). Acquired MS spectra were processed with the MaxQuant software suite V1.5.1.0^51, 52^ integrated with an Andromeda^53^ search engine. Database search was performed against a target-decoy database of *B. subtilis* (NCBI Assembly accession: ASM205596v1; GCA_002055965.1) containing 4,333 protein entries, and including additional 245 commonly observed laboratory contaminant proteins. Endoprotease Trypsin/P was set as the protease with a maximum missed cleavage of two. Carbamidomethylation (Cys) was set as a fixed modification. Label free quantification was enabled with a minimum ratio count of two. A false discovery rate of 10% was applied at the peptide and protein level individually for filtering identifications. Initial mass tolerance was set to 20 ppm. Intensity based absolute quantitation (iBAQ) option was enabled, but with log fit turned off. All other parameters were maintained as in the default settings. Finally, we obtained iBAQ values for 2,915 proteins at 10% false discovery rate.

### Transcriptome data analyses

Out of 4,333 protein coding genes with mapped reads we analysed 4,316 which passed the phylostratigraphic procedure (see below). First, we normalized raw counts of these 4,316 protein coding genes by calculating the fraction of transcripts (τ)^54^. This measure of relative expression, if multiplied with 10^6^, gives the widely used transcripts per million (TPM)^54^ quantity. As multiplication with the constant 10^6^ only serves to make values more human intuitive and does not influence further analysis in any way, we omitted the multiplication step and worked directly with τ. Given that this normalization allows cross-sample comparison and retains native expression variability^55^, a property crucial for correct estimation of partial concentrations in calculating phylo-transcriptomic measures like TAI^2^, we chose this approach as the most suitable for downstream calculations of evolutionary measures. After this normalization step, replicates were resolved by calculating their median, whereby we omitted replicates that had zero raw values. These transcriptome expression values were than used in all analyses except for assessing differential expression (see below). While preparing the transcriptome dataset for RNA expression profile correlations, visualization and clustering, we discarded the genes which had zero expression values in more than one stage, reducing thus the dataset to 4,296 genes. If a gene had only one stage with a zero-expression value, we imputed the zero-value by interpolating the mean of the two neighbouring stages (2 genes). In the case a zero-expression value was present in the first or the last stage of the biofilm ontogeny, we directly assigned the value of the only neighbour to it (134 genes in total). We decided on this procedure primarily with the aim to avoid erratic patterns in the visualization and clustering of mRNA expression profiles. To bring the gene expression profiles to the same scale, for every gene we performed the normalization to median and log2 transformed the obtained values. This procedure yielded per-gene normalized expression values for 4,296 genes (standardized expressions) which were visualized using the R *ggplot2* package V3.1.0 (Supplementary Information 3) and clustered with DP_GP_cluster^56^ with the maximum Gibbs sampling iterations set to 500 (Supplementary Information 1). For transcription factor and sigma factor expression profiles (Fig. 2, Extended Data Fig. 9) we selected genes which are regulating ≥ 10 operons based on the DBTBS database^57^. Biofilm-important genes (Fig. 2, Extended Data Fig. 7) and cell-to-cell signalling genes (Fig. 2, Extended Data Fig. 9) used for profile visualization were selected following relevant reviews^17–19^. Protein phosphatase and protein kinase genes (Fig. 2, Extended Data Fig. 9) were selected for profile visualization following the SubtiWiki database annotations^29^. The statistical significance of difference between average standardized expressions shown in Figure 2 was assessed by repeated measures ANOVA. To determine the similarity of transcriptomes across stages of biofilm ontogeny, we used Pearson’s correlation coefficients (R) calculated in an all-against-all manner and visualized in a heat map (Fig. 1b). Pairwise differential gene expression between individual stages of biofilm ontogeny (Extended Data Fig. 2) was estimated using a procedure implemented in the *DESeq2* V1.18.1 package. Raw counts of 4,316 protein coding genes were used as an input. Using the likelihood ratio test implemented in the same package, we also tested the overall differential expression for every gene across all stages of the biofilm ontogeny (Supplementary Table 5).

### Proteome data analyses

Out of 2,915 quantified proteins we analysed 2,907 which passed the phylostratigraphic procedure (see below). First, we calculated partial concentrations by dividing every iBAQ value by sum of all iBAQ values in the sample. After this normalization step replicates were resolved by calculating their median whereby we omitted replicates that had zero iBAQ values. This yielded normalized protein expression values that were used for calculating evolutionary indices. In preparing the proteome dataset for protein expression profile correlation, visualization and clustering, we further discarded genes which had zero-expression values in more than one stage, reducing thus the dataset to 2,543 proteins. If a protein had only one stage with a zero-expression value, we interpolated it by taking the mean of the two neighbouring stages (134 genes). In the case a zero-expression value was present in the first or the last stage of the biofilm ontogeny, we directly assigned it the value of the only neighbour (355 genes). To assess correlations between normalized transcriptome and proteome expression values, we calculated the Pearson’s correlation coefficient (R) on the matching 2,543 genes/proteins (Extended Data Fig. 6h, i). To bring the protein expression profiles to the same scale, for every gene we performed the normalization to median and log2 transformed the obtained values. This procedure yielded per-gene normalized expression values for 2,543 proteins (standardized expressions). Expression similarity across timepoints and replicates for 2,907 proteins was assessed using principal component analysis in R V3.4.2^45^ *stats* package. PCA plot was visualized using the R package *ggplot2* V3.1.0^49^ (Extended Data Fig. 1b).

### Enrichment analysis

Due to the poor gene annotations of our focal *B. subtilis* strain, we transferred annotations from *Bacillus subtilis* subsp. *subtilis* str. 168 (NCBI Assembly accession: ASM904v1; GCA_000009045.1) by establishing orthologs between the two strains (Supplementary Table 7). This was performed by calculating their reciprocal best hit using the blastp algorithm V2.7.1+^58^ with 10^-5^ e-value cut-off. Functional annotations for the *B. subtilis* 168 strain were retrieved from the SubtiWiki database^29^ (October 23, 2019). Functional enrichment of transcriptome and proteome clusters, and individual biofilm timepoints was estimated using the one-way hypergeometric test. For each timepoint genes that had expression in that timepoint 0.5 times (log2 scale) above the median of their overall expression profile were tested for functional enrichment. *P* values were adjusted for multiple comparisons using the Yekutieli and Benjamini procedure^59^.

### Evolutionary measures

The phylostratigraphic procedure was performed as previously described^2, 25^. Following the recent phylogenetic literature^60–73^, we constructed a consensus phylogeny that covers divergence from the last common ancestor of cellular organisms to the *B. subtilis* as a focal organism (Extended Data Fig. 3; Supplementary Information 2). We chose the nodes based on their phylogenetic support in the literature^60–73^, importance of evolutionary transitions and availability of reference genomes for terminal taxa. The full set of protein sequences for 926 terminal taxa were retrieved from ENSEMBL^74^ (922) and NCBI (4) databases. After checking consistencies of the files, leaving only the longest splicing variant per gene for eukaryotic organisms and adding taxon tags to the sequence headers of all sequences, we prepared a protein sequence database for sequence similarity searches (Supplementary Table 2). To construct the phylostratigraphic map^2, 25^ of *B. subtilis*, we compared 4,333 *B. subtilis* proteins to the protein sequence database by blastp algorithm V2.7.1+ with the 10^-3^ e-value threshold. After discarding all protein sequences which did not return their own sequence as a match 4,317 protein sequences left in the sample. These 4,317 protein sequences were than mapped on the 12 internodes (phylostrata) of the consensus phylogeny using a custom-made pipeline (Supplementary Table 2). We assigned a protein to the oldest phylostratum on the phylogeny where a protein still had a match (Dollo’s parsimony)^2, 25^. Using expression values for 4,316 protein coding genes we calculated the transcriptome age index (TAI), i.e. the weighted mean of phylogenetic ranks (phylostrata), for each ontogenetic stage (Fig. 1e). To analyse the proteome data in similar fashion, we introduced the proteome age index (PAI):

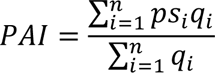

where ps*i* is an integer which represents the phylostratum of the protein *i*, *q*i is iBAQ value of the protein *i* that acts as weight factor and *n* is the total number of proteins analysed. In total, we used 2,907 proteins for PAI calculation. To estimate evolutionary divergence rates of *B. subtilis* proteins, we found 3,094 orthologs in *Bacillus licheniformis* str. DSM 13 (NCBI Assembly accession: ASM1164v1; GCA_000011645.1) by reciprocal best hits using blastp with 10^-5^ e-value threshold (Supplementary Table 3). Of the available *Bacillus* species, *B. licheniformis* provides the best balance between number of detected orthologues and evolutionary distance of species pair for calculating evolutionary rates. We globally aligned *B. subtilis* – *B. licheniformis* orthologous pairs using the Needleman–Wunsch algorithm and then constructed codon alignments in pal2nal^75^. The non-synonymous substitution rate (dN) and the synonymous substitution rate (dS) were calculated using the Comeron’s method^76^. The whole procedure of obtaining dN and dS was performed in the R package *orthologr* V0.3.0.9000^77^. Using dN values of 3,091 genes, we calculated the transcriptome nonsynonymous divergence index (TdNI; Fig. 1f), *i.e.* the weighted arithmetic mean of nonsynonymous gene divergence:

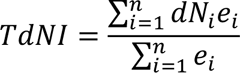

where dN*i* is a real number which represents the nonsynonymous divergence of gene *i*, *e*i is the transcriptome expression value of the gene *i* that acts as weight factor and *n* is the total number of genes analysed. Using dS values of 2,212 genes, we calculated transcriptome synonymous divergence index (TdSI; Extended Data Fig. 5a), *i.e.* the weighted arithmetic mean of synonymous gene divergence:

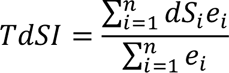

where dS*i* is a real number which represents the synonymous divergence of gene *i*, *e*i is the transcriptome expression value of the gene *i* that acts as weight factor and *n* is the total number of genes analysed. To analyse proteome data in similar fashion, we introduced the PdNI and PdSI. Using dN of 2,329 proteins, we calculated the proteome nonsynonymous divergence index (PdNI; Fig. 1h), i.e. the weighted arithmetic mean of nonsynonymous divergence:

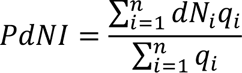

where dN*i* is a real number which represents the nonsynonymous divergence of protein *i*, *q*i is the normalized protein expression value of the protein *i* that acts as weight factor and *n* is the total number of proteins analysed. Using expression values of 1,755 proteins, we calculated proteome synonymous divergence index (PdSI; Extended Data Fig. 5b), i.e the weighted arithmetic mean of synonymous divergence:

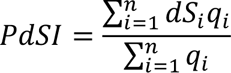

where dS*i* represents the synonymous divergence value of protein *i*, *q*i is the normalized protein expression value of protein *i* that acts as weight factor and *n* is the total number of proteins analysed. Using 4,316 transcriptome expression values and “measure independent of length and composition” (MILC) ^79^ values we calculated transcriptome codon usage bias index (TCBI), *i.e.* the weighted arithmetic mean of codon usage bias:

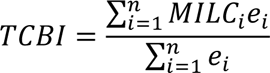

where *MILC* is a real number which represents the codon usage bias of gene *i*, *e*i is the transcriptome expression value of the gene *i* that acts as weight factor and *n* is the total number of genes analysed. Using 2,907 protein expression and MILC values we calculated proteome codon usage bias index (PCBI), i.e. the weighted arithmetic mean of codon usage bias:

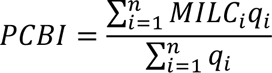

where *MILC* is a real number which represents the codon usage bias of protein *i*, *q*i is the normalized protein expression value of protein *i* that acts as weight factor and *n* is the total number of proteins analysed. MILC values were obtained from R package coRdon V1.3.0^80^, with respect to codon usage bias of ribosomal genes. The whole procedure of obtaining TdNI, TdSI, PdNI and PdSI was made in R package *orthologr* V0.3.0.9000. The statistical analysis for TAI, PAI, TdNI, TdSI, PdNI, PdSI, TCBI, PCBI was calculated using the R package *myTAI* V0.9.0^78^.

### Imaging

Images of biofilms at sampling timepoints (Fig. 1a) along with the time-lapse video (Supplementary Video 1) were taken using the Sony Alpha a7 II mirrorless camera attached to a Zeiss Stemi 2000-C stereo-microscope with a NEX/T2 adapter (Novoflex) at 30 °C and average relative humidity of 74.5% using the automatic camera settings. The time-lapse video was produced from 1,345 shots taken during 14 days in 15 min intervals using the Adobe After Effects CC 2017 software at 24 fps.

## Data Availability

All transcriptome data have been deposited in NCBI’s Gene Expression Omnibus and are accessible through GEO Series accession number GSE141305. All mass spectrometry proteomics data have been deposited in the PRIDE database and are accessible through the dataset identifier PXD016656.

---

**Extended Data Fig. 1.**
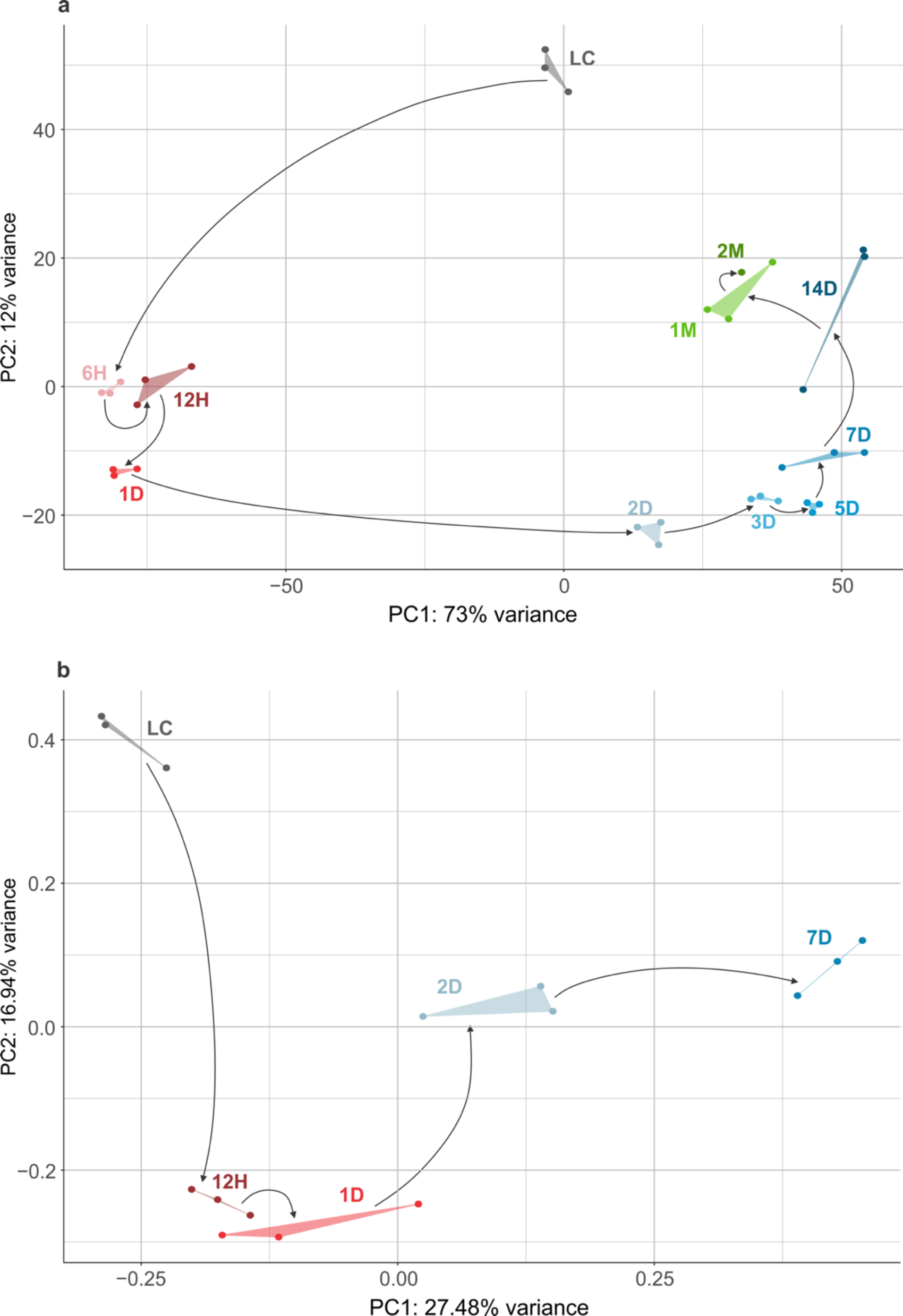
Principal component analysis (PCA) of transcriptomes and proteomes shows a punctuated organization of biofilm growth. PCA of a, transcriptome and b, proteome data (see Methods). Biofilm growth timepoints (LC, 6H, 12H, 1D, 2D, 3D, 5D, 7D, 14D, 1M and 2M) are shown in different colours, where grey represents the liquid culture (LC), different shades of red early (6H-1D), blue mid (2D-14D) and green late (1M-2M) biofilm period. Replicates are in the same colour and connected with lines. Black arrows correspond to the experimental timeline of biofilm growth that starts with LC and ends at 2M.

**Extended Data Fig. 2.**
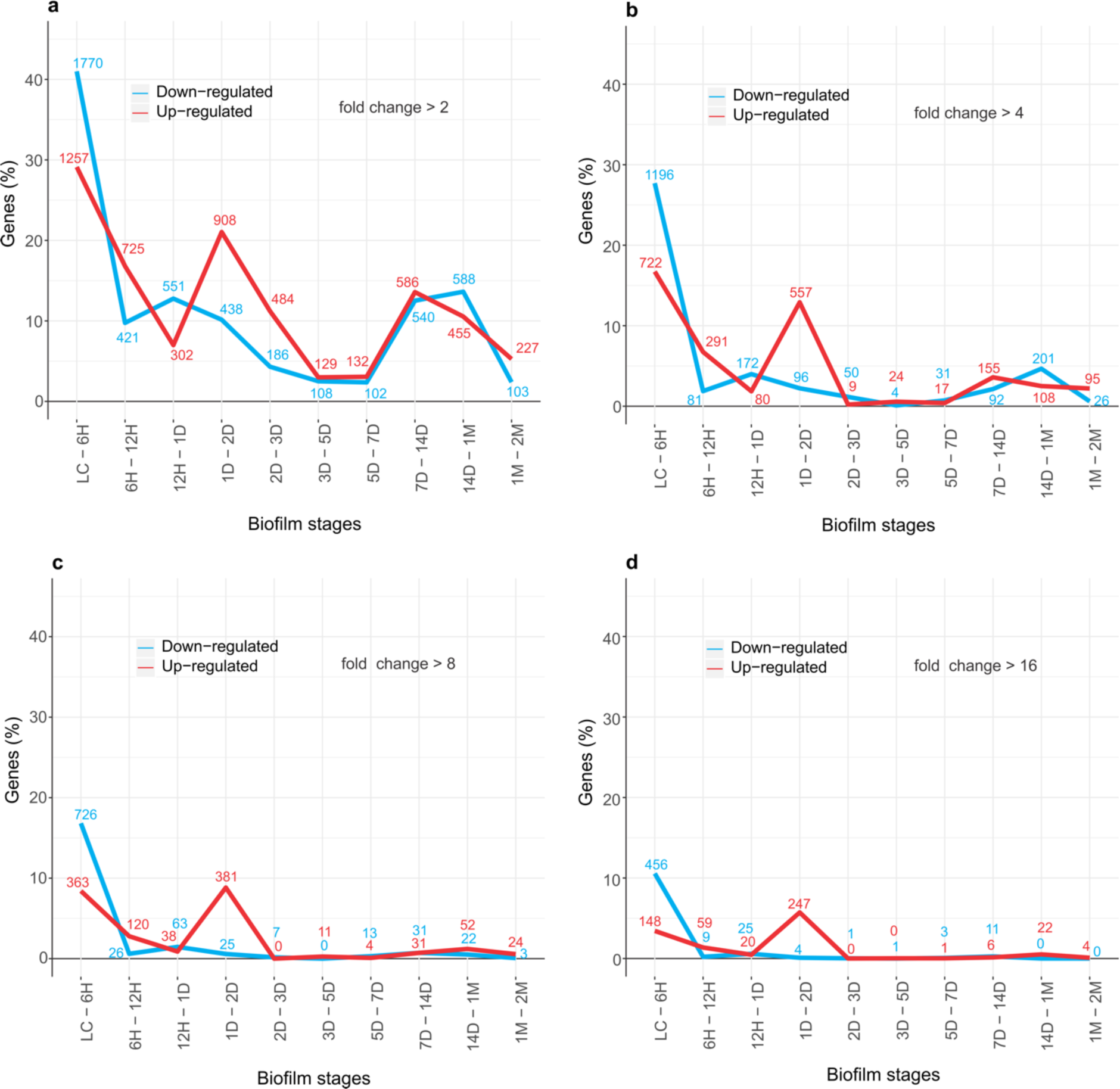
Differential transcription in pairwise comparisons between successive biofilm timepoints. Bursts of differentially transcribed genes are detected at inoculation (LC-6H), 1D-2D and at 7D-14D transitions. The numbers above lines indicate the number of differentially expressed genes. Down-regulated genes are in blue, up-regulated are in red. a, fold change cut-off >2; b, fold change cut-off >4; c, fold change cut-off >8 and d, fold change cut-off >16. The highest number of highly upregulated genes are at the 1D-2D transition (fold change>8 and >16). Differentially expressed genes were determined by DeSeq2 with *p* value (adjusted for multiple comparisons) cut-off set at 0.05.

**Extended Data Fig. 3.**
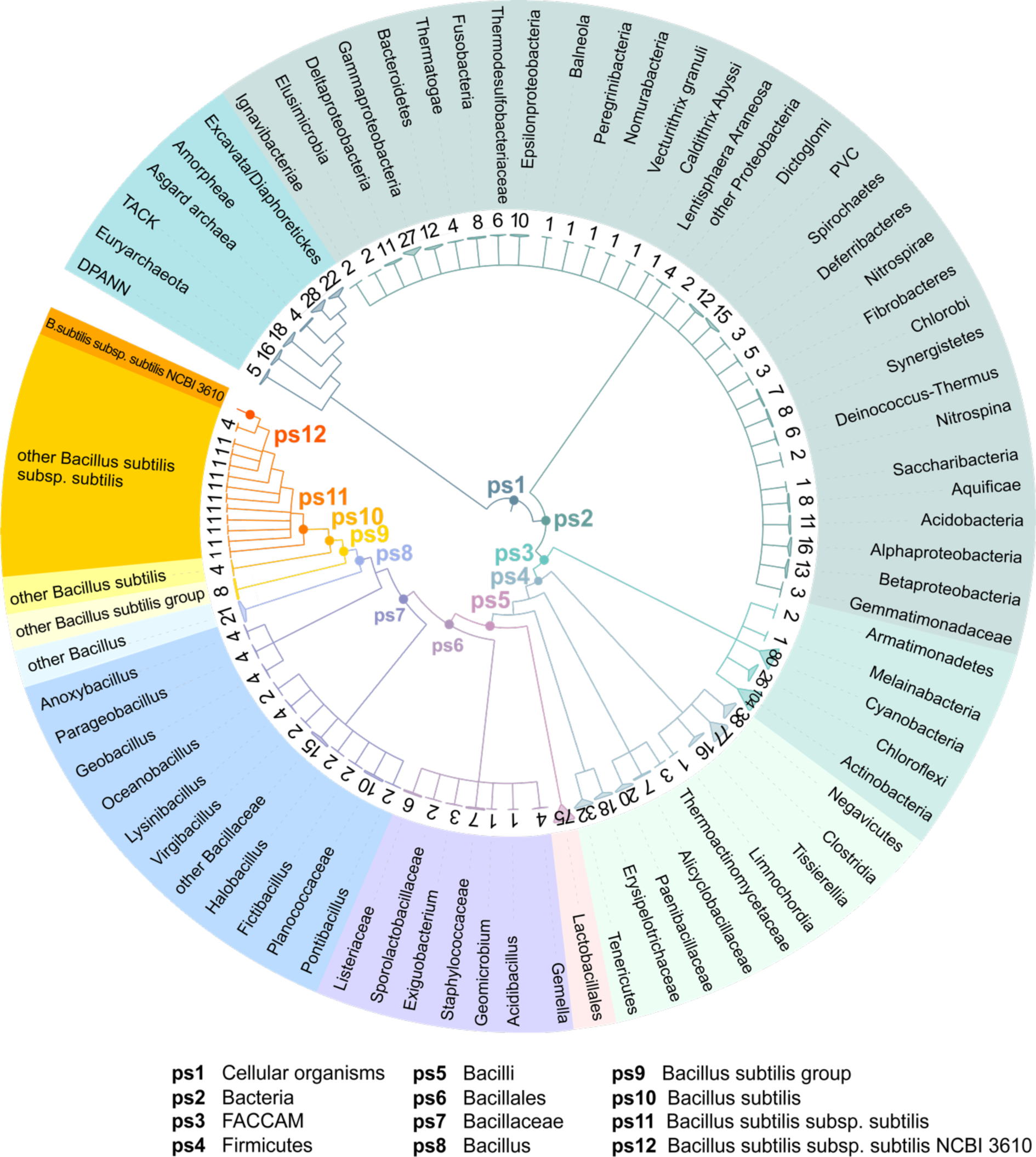
The consensus phylogeny used in the phylostratigraphic analysis. The consensus tree covers divergence from the last common ancestor of cellular organisms to *Bacillus subtilis* str. NCBI 3610 as a focal organism (see Supplementary Information 2 for fully resolved tree) and is constructed based on the relevant phylogenetic literature^60–73^, importance of evolutionary transitions and availability of reference genomes. Twelve internodes (phylostrata) considered in the phylostratigraphic analysis are marked by ps1-ps12. The numbers at the top of terminal nodes represent the number of species in the fully resolved tree and correspond to the genomes used to populate the reference database for sequence similarity searches. The abbreviation FACCAM (ps3) stands for Firmicutes, Actinobacteria, Chloroflexi, Cyanobacteria, Armatimonadates and Melainabacteria.

**Extended Data Fig. 4.**
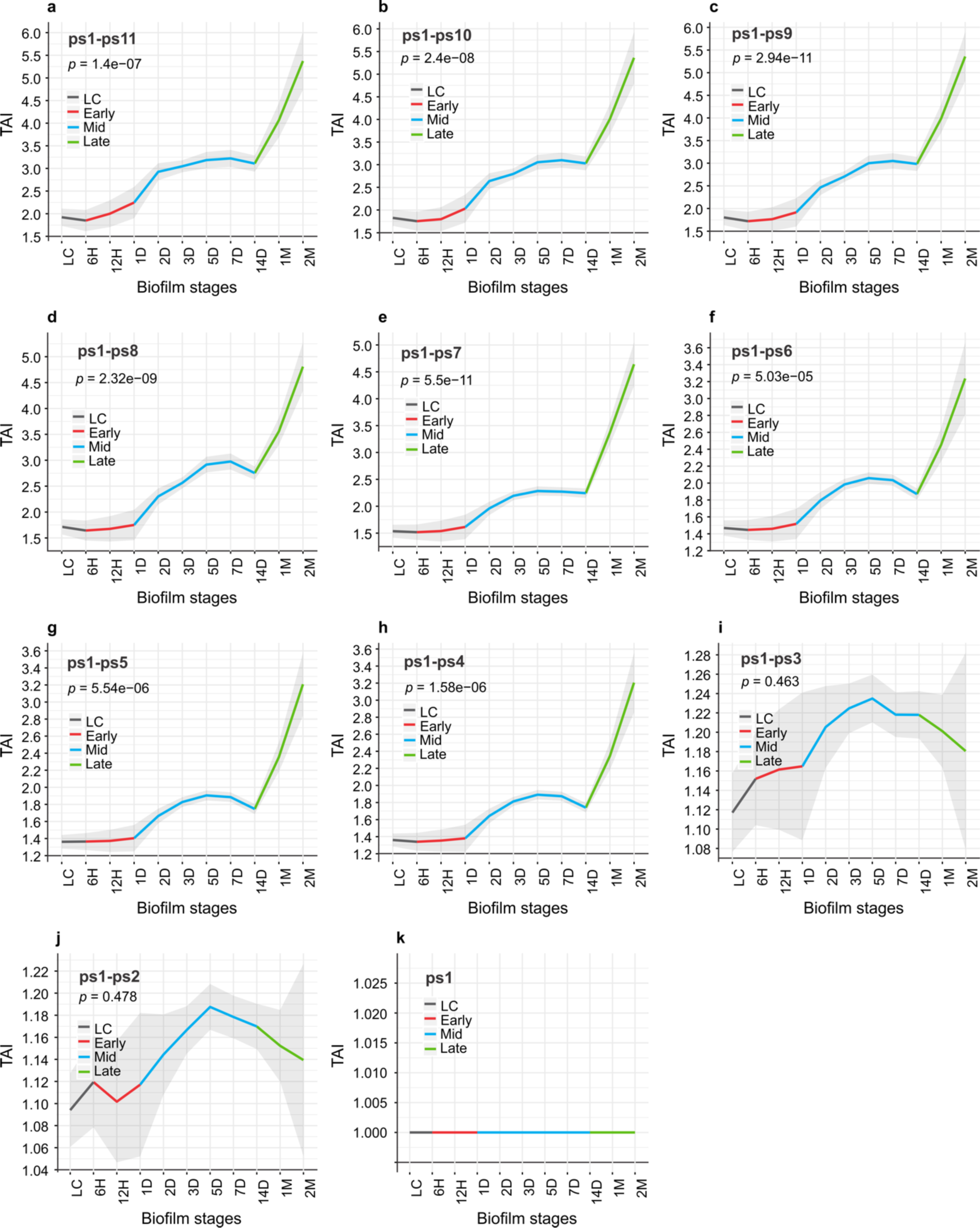
Recapitulation pattern is significant from the origin of Firmicutes at ps4. Transcriptome age index (TAI) was calculated on the reduced datasets by progressively removing genes from younger phylostrata a, ps1-ps11 (n = 4,293); b, ps1-ps10 (n = 4,218); c, ps1-ps9 (n = 4,186); d, ps1-ps8 (n = 4,097); e, ps1-ps7 (n = 3,923); f, ps1-ps6 (n = 3,778); g, ps1-ps5 (n = 3,707); h, ps1-ps4 (n = 3,683); i, ps1-ps3 (n = 3,239); j, ps1-ps2 (n = 3,159) and k, ps1 (n = 2,562). Depicted *p* values are obtained by the flat line test and grey shaded areas represent ± one standard deviation estimated by permutation analysis (see Methods). Early (red), mid (blue) and late (green) periods of biofilm growth are colour coded.

**Extended Data Fig. 5.**
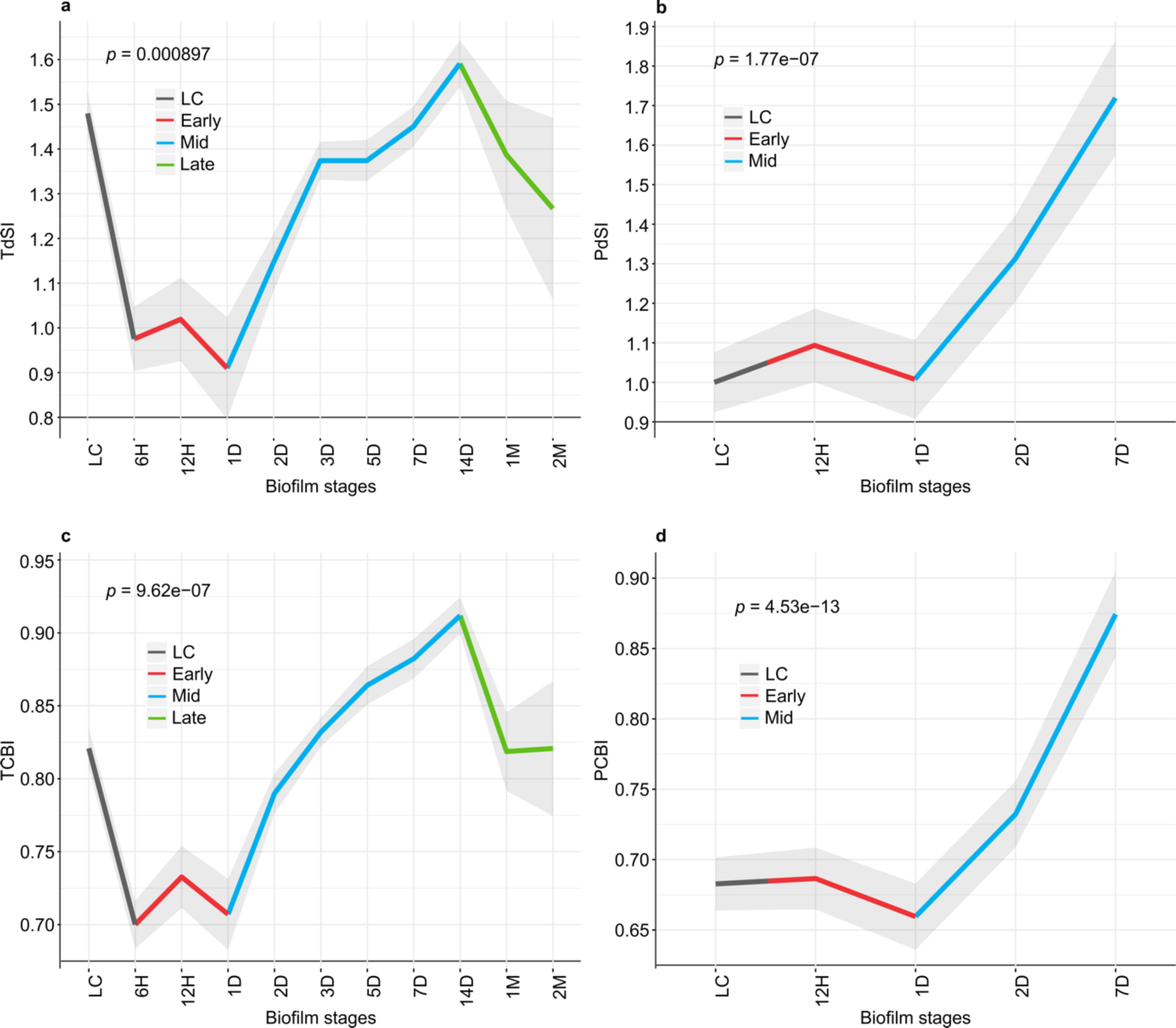
Synonymous divergence and codon usage bias are co-dependent. a, Transcriptome synonymous divergence index (TdSI) shows resemblance to c, transcriptome codon bias index (TCBI). b, Proteome synonymous divergence index (PdSI) shows resemblance to d, proteome codon bias index (PCBI). Depicted *p* values are obtained by the flat line test and grey shaded areas represent ± one standard deviation estimated by permutation analysis (see Methods). Early (red), mid (blue) and late (green) periods of biofilm growth are colour coded.

**Extended Data Fig. 6.**
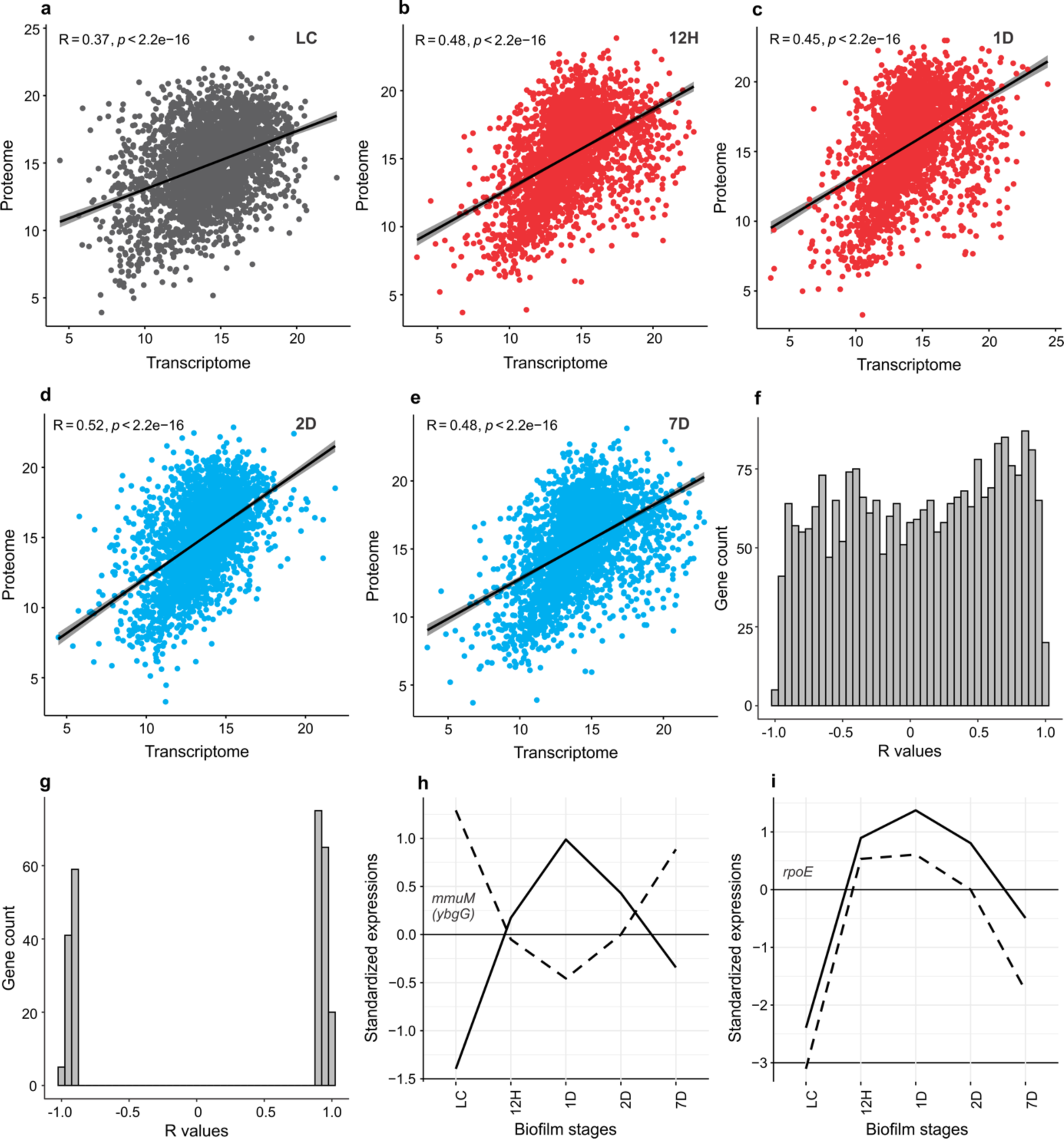
Transcriptome and proteome data show relatively low correlation. Correlation between transcriptome and proteome normalized expression values on -log2 scale calculated for each biofilm timepoint separately. Pearson’s correlation coefficients (R) with corresponding *p* values are shown. Genes that had zero expression values in either proteome or transcriptome were excluded. a, LC (n = 2,720 genes); b, 12H (n = 2,763 genes); c, 1D (n = 2,721 genes); d, 2D (n = 2,590 genes) and e, 7D (n = 2,318 genes). f, Distribution of Pearson’s correlation coefficients (R) calculated for every gene between its transcriptome and proteome standardized expression profile across biofilm ontogeny (n = 2,543 genes). g, Distribution of only significant Pearson’s correlation coefficients (R) (*p* < 0.05, n = 265 genes) from f. Significant negative correlation show 105 (4.1%) and significant positive 160 (6.3%) genes. h, An example of gene (*mmuM*; *ybgG B. subtilis* 168 strain name) that shows significant negative Pearson’s correlation (*p* = 0.009, R = -0.96) between the transcriptome (dashed line) and the proteome (solid line). i, An example of gene (*rpoE*) that shows significant positive Pearson’s correlation (*p* = 0.003, R = 0.98) between the transcriptome (dashed line) and the proteome (solid line).

**Extended Data Fig. 7.**
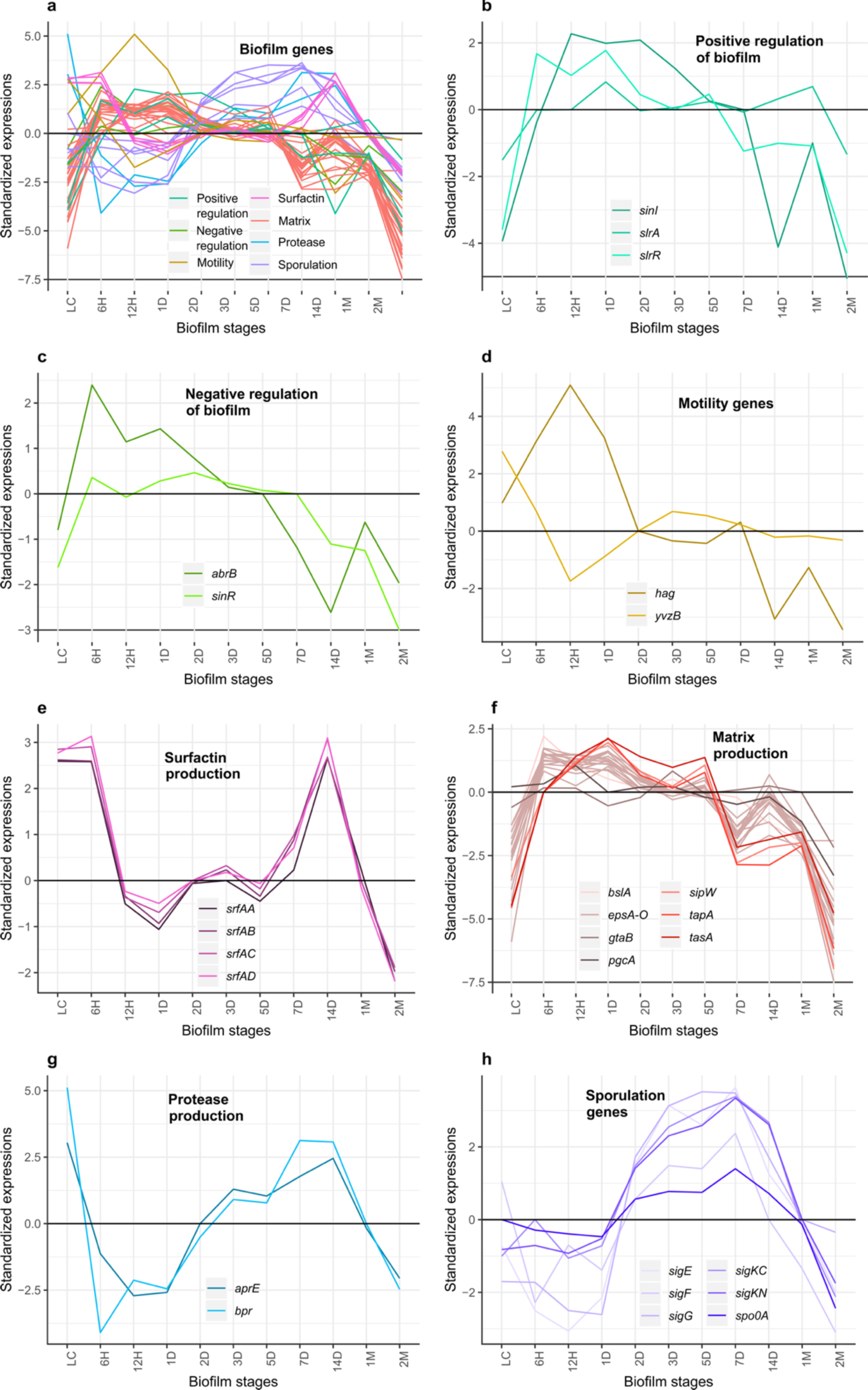
Standardized transcription profiles of key biofilm genes. a, All considered biofilm genes grouped in seven categories (n = 40); b, Positive regulators of biofilm growth (n = 3); c, Negative regulators of biofilm growth (n = 2); d, Biofilm important cell motility genes (n = 2); e, Surfactin genes (n = 4); f, Matrix genes(n = 21); g, Biofilm important proteases (n = 2); h, Sporulation genes (n = 6). Black horizontal line represents the median of standardized transcriptome expression values. See Supplementary Table 6 for full list of genes used for profile visualization, as well as their corresponding values.

**Extended Data Fig. 8.**
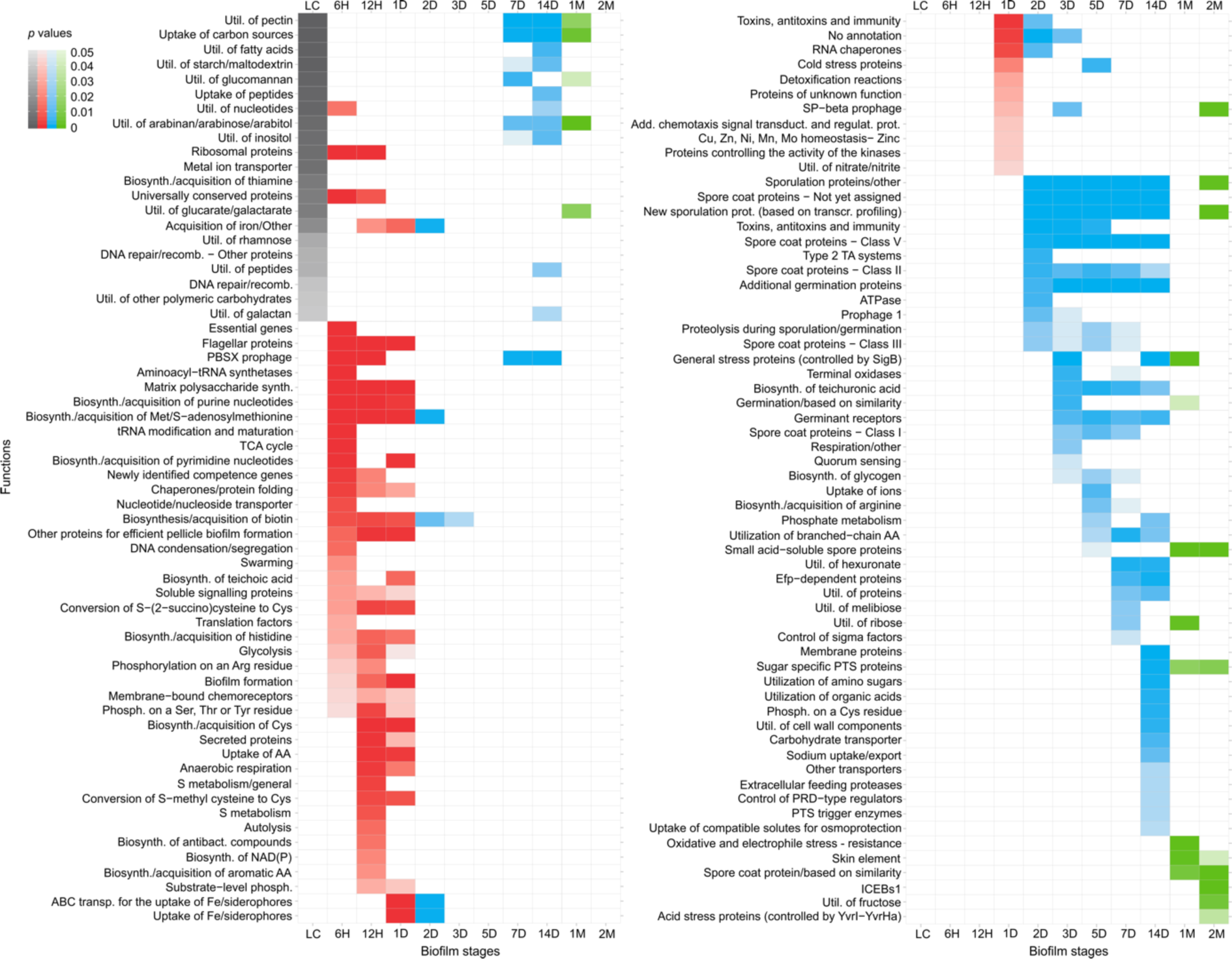
Biofilm ontogeny is a punctuated process organized in functionally discreate stages. Enrichment analysis of SubtiWiki functional categories (maximal ontology depth) in a respective biofilm growth timepoint for genes with transcript expression 0.5 times (log2 scale) above the median of their overall transcription profile. Similar results are obtained for other transcription level cut-offs and SubtiWiki functional annotation ontology depths (see Supplementary Table 4). Colouring follows biofilm growth periods: LC (grey), early (red), mid (blue), late (green). Functional enrichment is tested by one-tailed hypergeometric test and *p* values are adjusted for multiple testing (see Methods).

**Extended Data Fig. 9.**
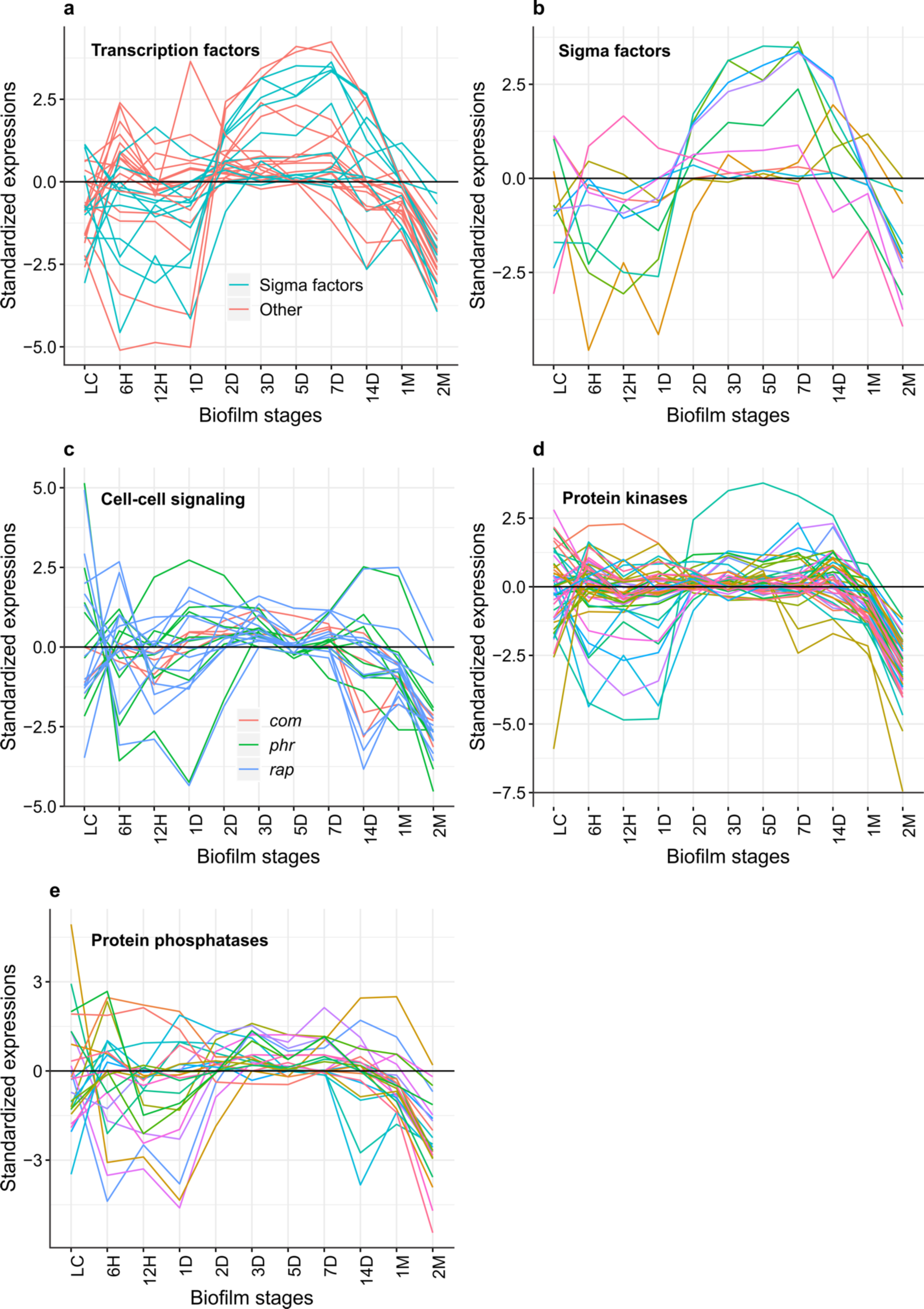
Multicellularity-important genes show the strongest transcriptome expression in the mid-period of biofilm growth. Standardized transcription profiles: a, Transcription factors (n = 28); b, Sigma factors (n = 11); c, Cell to cell signalling genes (n = 24); d, Protein kinases (n = 49); e, Protein phosphatases (n = 24); f, Key biofilm genes (n = 40). Black horizontal line represents the median of standardized transcriptome expression values. For individual values and profiles of all considered genes see Supplementary Table 6 and Supplementary Information 3.

**Extended Data Fig. 10.**
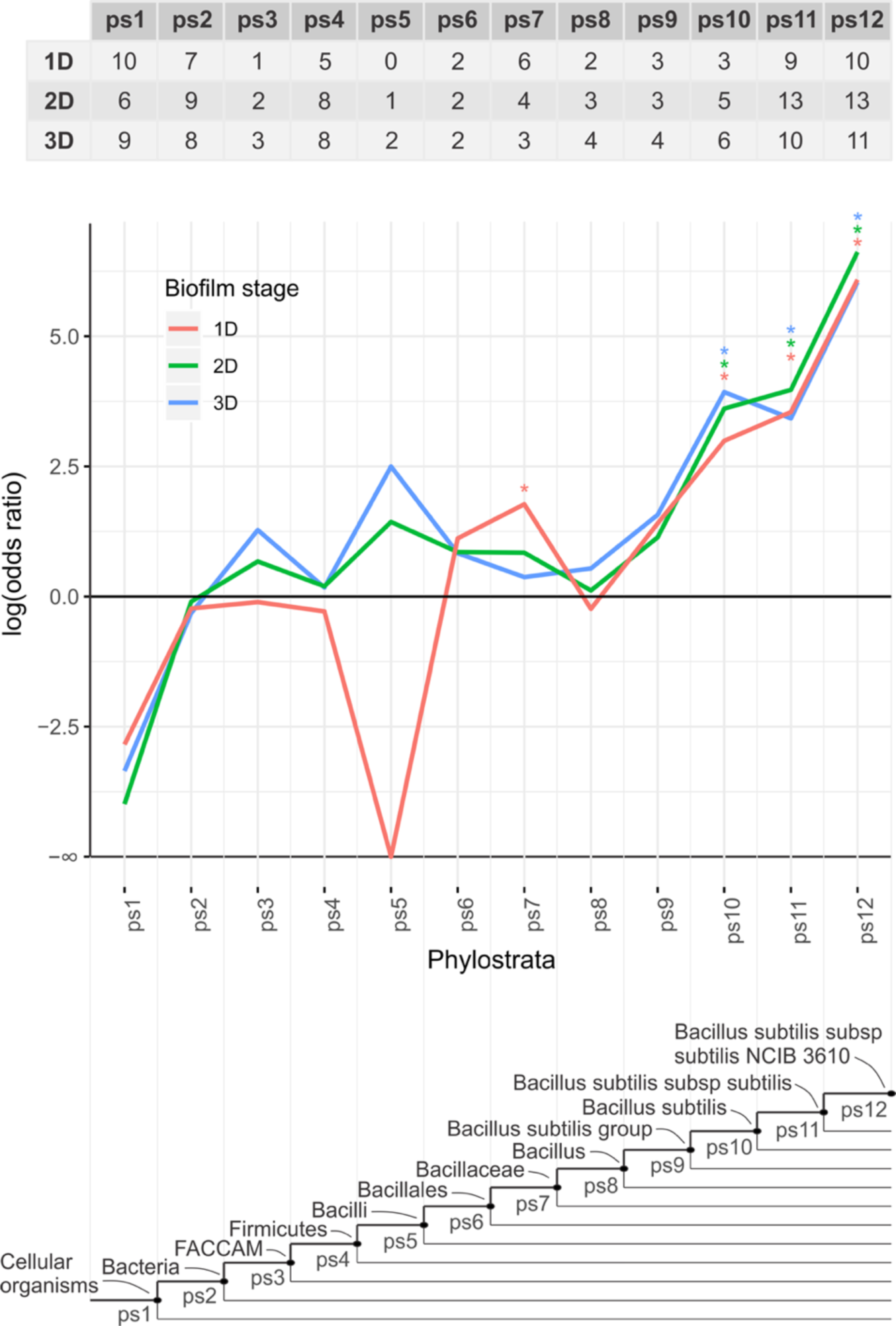
Distribution of functionally unannotated genes on the phylostratigraphic map. Evolutionary origin of genes that contribute to the enrichment of the functional term “no annotation” at 1D, 2D and 3D timepoints (see Fig. 3). The table shows the number of genes without functional annotation per phylostratum for 1D, 2D and 3D timepoints. Phylostratigraphic enrichment is tested by one-tailed hypergeometric test and *p* values are adjusted for multiple testing (* *p* < 0.05). The abbreviation FACCAM (ps3) stands for Firmicutes, Actinobacteria, Chloroflexi, Cyanobacteria, Armatimonadates and Melainabacteria.

